# Parallel clines and sympatric divergence in ecological and mating traits shape speciation in Lake Victoria cichlids

**DOI:** 10.1101/2024.10.31.621340

**Authors:** Grégoire Gautier, Anna Mahulu, Ole Seehausen, Pooja Singh

## Abstract

Speciation often emerges from the complex interplay of geography, demography, ecology, and sexual selection, yet disentangling their relative contributions remains a challenge. In this study, we examine two species complexes of rock-dwelling Lake Victoria cichlids in the genera *Pundamilia* and *Neochromis* that are distributed along a continuous geographic gradient of rocky shores. Using morphometric analyses and male nuptial coloration data, we reveal clinal craniofacial divergence following an isolation by distance model of evolution, punctuated by strong local effects. While the *Pundamilia* cline exhibited modular shifts in craniofacial anatomy linked to diet-specific morphology, variation in the *Neochromis* cline was dominated by allometric constraints, suggesting different evolutionary processes at play. At the single site where the terminal populations of both clines co-occur, each pair of sister species showed pronounced divergence in morphology and male nuptial coloration, consistent with ecological and sexual character displacement facilitating co-existence. Our findings highlight that clinal morphological variation in parapatry and more abrupt shifts in sympatry interact to generate and maintain species. These rocky shore cichlids provide a rare opportunity to investigate the speciation continuum in one of the world’s youngest adaptive radiations.

## INTRODUCTION

Speciation is a fundamental evolutionary process through which populations diverge to the extent that they can coexist as distinct species. While genetic patterns in speciation are increasingly well documented, the integration of biogeographic and ecological mechanisms is less well studied, particularly in young species (Losos and Glor, 2003; Lyu *et al*., 2015; Stankowski *et al*., 2024). In gradually distributed systems, classical theory predicts that gene flow between populations decreases with geographic distance, leading to the gradual accumulation of genetic, and potentially phenotypic differences across space—a process known as isolation by distance (IBD; Wright, 1943). The consequence of IBD may be that populations diverge in a cline-like fashion and if the cline is sufficiently long and steep, populations at opposite ends may become sufficiently differentiated to constitute two distinct species. However, the stable coexistence of such terminal species in sympatry typically requires additional processes, most notably ecological character displacement (ECD), which reduce niche overlap and alleviate interspecific competition (Brown and Wilson, 1956; Goldberg and Lande, 2006; Weber and Strauss, 2016). Yet, few empirical studies have traced how IBD-driven divergence and community-level interactions jointly shape ecological and mating trait differentiation in spatially continuous systems.

Lake Victoria in East Africa hosts an eco-morphologically diverse adaptive radiation of hundreds of cichlid fishes that evolved in less than 16,000 years within species rich communities (Johnson *et al*., 1996; Seehausen, 1996; Sturmbauer *et al*., 2011). Both ecological factors—such as variation in diet and habitat use—and sexual selection via female mate choice and male-male competition contribute to speciation and trait divergence in this radiation (Bouton *et al*., 1999; Maan *et al*., 2004; Dijkstra *et al*., 2007). The geography of trait divergence, and the degree to which it is shaped by spatial isolation versus interactions in sympatry, has only been studied in a few cases (Magalhaes *et al*., 2010; Terai *et al*., 2006; Seehausen *et al*., 2008) and no study has yet addressed possible interactions between clinal divergence and secondary contact. Seehausen (1996) described two species complexes in the *Pundamilia* and *Neochromis* genera that have populations distributed along the southern rocky shores and islands of Lake Victoria (Fig. 1). Both these complexes display gradual ecomorphological changes and male nuptial coloration shifts across populations that extend around an open lake area with deep water, which acts as a dispersal barrier for these shallow water rocky shore specialised fishes. In both of these species complexes, the two most strongly differentiated populations occur in sympatry at one location (Juma island) where they form two distinct co-existing species that appear to be connected by clines of intermediate populations in parallel circuits around the opening of the Mwanza Gulf (Fig. 1b). Populations of these complexes may represent steps along a speciation continuum from geographically adjacent populations that exchange migrants regularly to two reproductively differentiated species, and offer a rare and powerful comparative framework for exploring how ecological adaptation and reproductive differentiation evolve during speciation across a shared geographic landscape.

**Figure 1:**
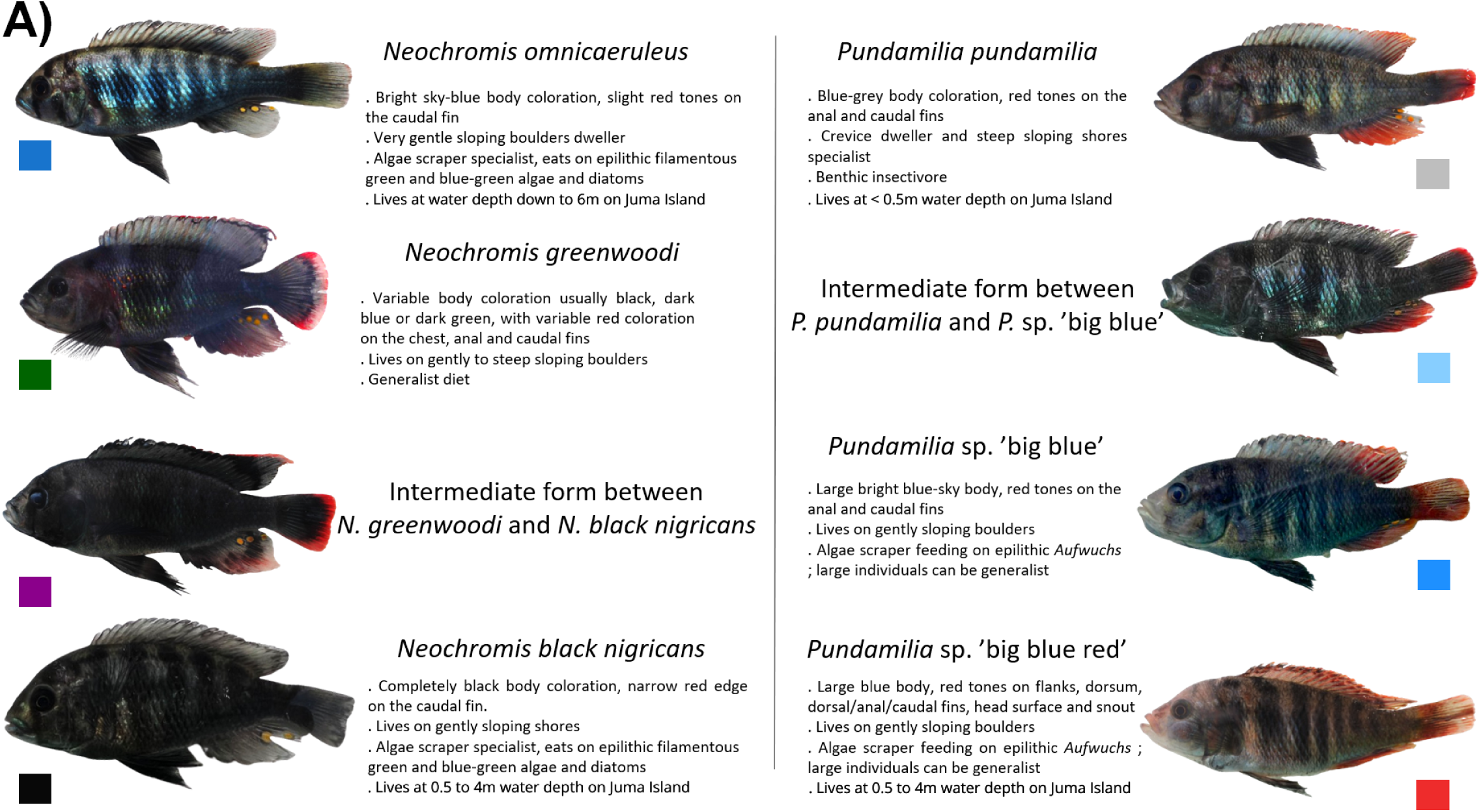
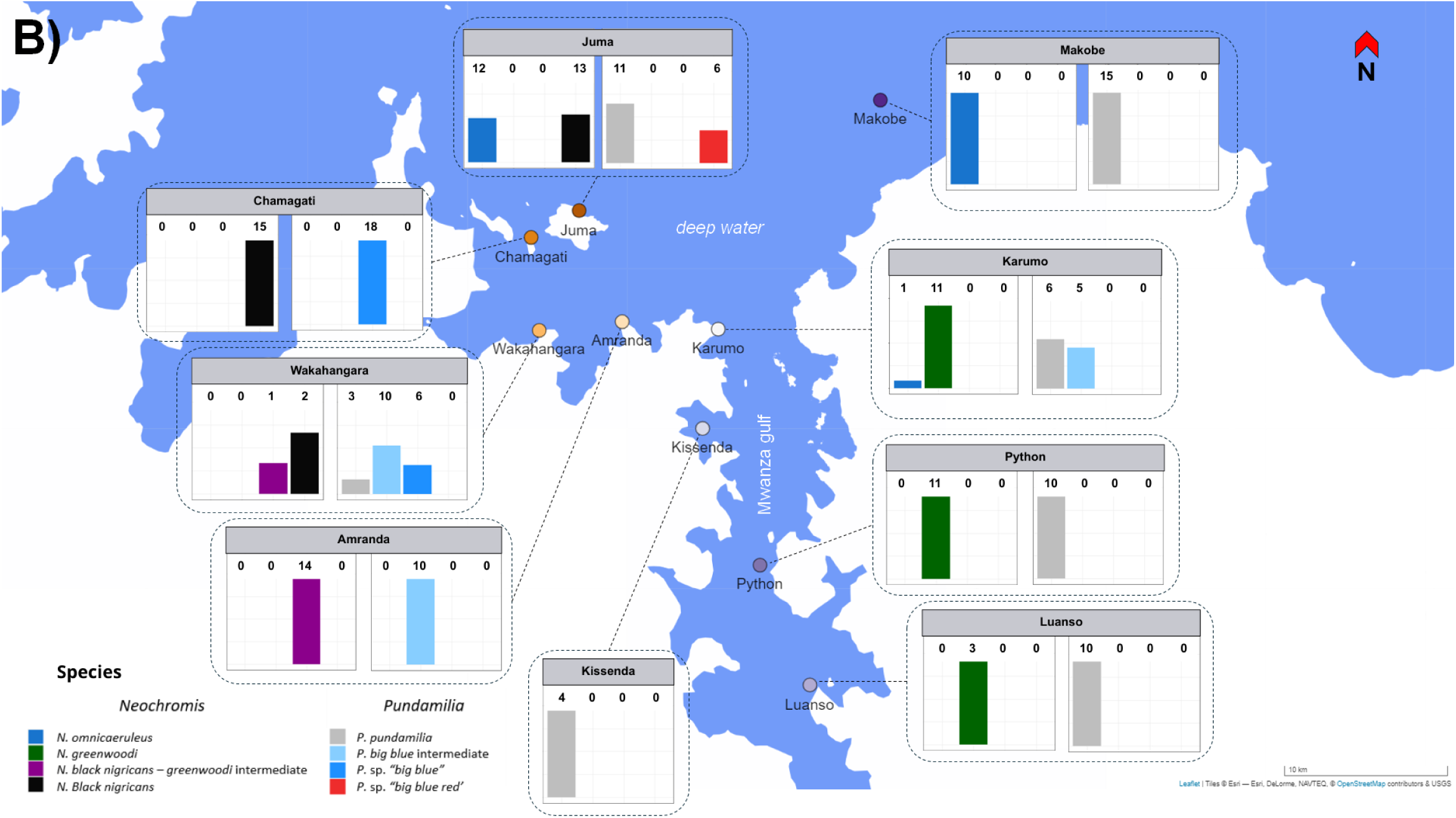
Description of the *Pundamilia* and *Neochromis* species complexes used for the study, and summary of the sampling in Southern Lake Victoria. A) Details of male nuptial coloration, habitat characteristics, and diet, based on Seehausen 1996. Note that male cichlid nuptial coloration fades rapidly upon capture. Photos therefore tend to show faded colours compared to *in situ* conditions. Colour coding of species and intermediates will be used consistently in the manuscript figures. **B)** Species composition at sampling sites in the Mwanza gulf and Sengerema region of Southern Lake Victoria, for *Pundamilia* (*n* = 111) and *Neochromis* (*n* = 92) complexes (left and right respectively; no fish from Kissenda island was used for the study in *Neochromis*). Numbers above the bars indicate the number of fish for each species at one location.

Here, we investigate patterns of ecological and mating trait divergence in *Pundamilia* and *Neochromis* populations distributed along the southern shores and islands of Lake Victoria. By examining craniofacial morphology and male nuptial coloration across multiple locations, our goal is to test predictions of trait divergence through IBD, and ecological and sexual character displacement in secondary sympatry. In doing so, we aim to contribute to a broader understanding of how neutral processes, ecological and sexual selection interact during speciation in parapatry.

## MATERIALS AND METHODS

### Fish sampling and linear measurements

Specimens analysed in this study were collected in 2022 by P. Singh, O. Seehausen, and A. Mahulu in Southern Lake Victoria around the Mwanza Gulf and Sengerema region (Tanzania). The Sengerema region comprises Kissenda, Karumo, Amranda, Wakahangara, Chamagati and Juma locations, while Python and Luanso Islands lie within the Southern Mwanza gulf (Fig. 1). Makobe island lies just outside the gulf approximately five kilometres north of the mainland. Additional specimens were included from previous fieldwork (1996 and 2014, O. Seehausen) to increase sample sizes for Makobe and Python Islands.

Sampling focused on rocky shore habitats along mainland and island coasts, following the distribution patterns described by Seehausen (1996) ensuring representative coverage of focal populations. Fish were collected using gillnets and angling and were sacrificed humanely using clove oil anesthetic in accordance with the Tanzanian and Swiss animal welfare regulations. Specimens were fixed in 4.0% formalin and later transferred to ethanol. For the *Pundamilia* complex, we obtained 111 male individuals belonging to the populations (or intermediate forms) of interest: *Pundamilia pundamilia*, *Pundamilia* sp. “big blue”, and *Pundamilia* sp. “big blue red”. For the *Neochromis* complex, we obtained 92 males from the populations of interest: *Neochromis omnicaeruleus*, *Neochromis greenwoodi*, and *Neochromis* sp. “*black nigricans*”. Only males were included in the analysis because females cannot readily be distinguished between the closely related species by eye and also because sexual dimorphism in size and morphology make the joint analysis difficult.

Linear measurements were conducted on twelve linear traits characterising head and body shape, and known to capture differences between haplochromine species and respond to feeding-related variation (Barel *et al*., 1976; Cooper *et al*., 2010; Hulsey *et al*., 2010; Hulsey and León, 2005; Magalhaes *et al*., 2010; Seehausen, 1996): body depth (BD), head length (HL), head width (HW), lower jaw length (LJL), lower jaw width (LJW), snout length (SnL), snout width (SnW), eye length (EyL), eye depth (EyD), cheek depth (ChD), pre-orbital depth (POD), and inter-orbital width (IOW). All measurements were taken using digital calipers following standard morphometric protocols for cichlids (Barel *et al*., 1976)(Fig. 2). Trait values were corrected for size using log–log regressions against standard length (SL)(Fleming and Gross, 1994; Magalhaes *et al*., 2010, 2009).

**Figure 2:**
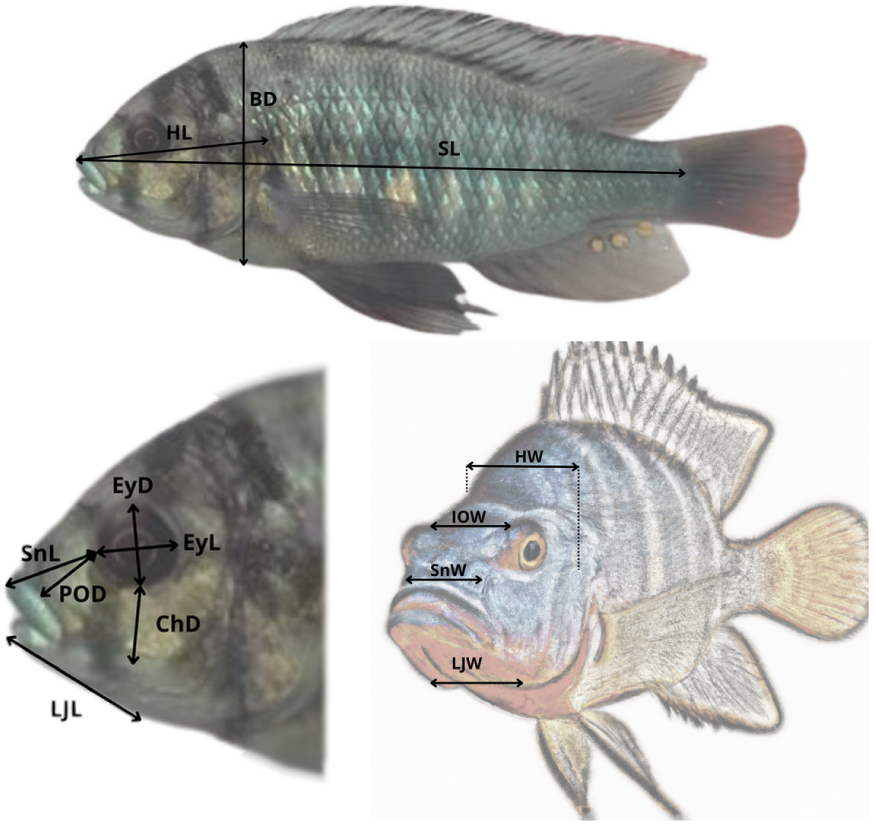
Linear measurements used for the morphological analysis of *Pundamilia* and *Neochromis* populations, based on Barel *et al*. 1976. SL: Standard length; BD: Body depth; HL: Head length; EyL: Eye length; EyD: Eye depth; ChD: Cheek depth; POD: Pre-orbital depth; SnL: Snout length; LJL: Lower jaw length; SnW: Snout width; LJW: Lower jaw width; IOW: Inter-orbital width; HW: Head width.

### Statistical analyses

All analyses were performed in R v4.4.3 (R Core Team, 2025) using RStudio 2023.12.1 (Posit, 2024). *Pundamilia* and *Neochromis* were analysed separately but with an identical workflow. R script for all analyses is provided as a supplementary file (File S1).

#### Craniofacial morphology variation

We used principal component analysis (PCA; *FactoMineR*: Husson *et al*., 2024; *factoextra*: Kassambara and Mundt, 2020) on size-corrected trait values to quantify morphological variation across three datasets: (1) all individuals, (2) individuals excluding Juma Island’s *N. omnicaeruleus* and *P.* sp. “big blue red”, and (3) sister species in sympatry. Dataset (2) aimed to capture potential continuous morphological variation in parapatric populations, while excluding populations thought to represent a distinct evolutionary origin. To identify functional modules, we combined two approaches: (i) hierarchical clustering of traits to detect correlated groups, and (ii) inspection of trait loadings on the first four PCA axes, selected using the elbow method. Finally, to visualize spatial structuring of morphology, we plotted mean PC scores per location for dataset (1).

#### Isolation by distance along distribution

We set four alternative spatial hypotheses of linear connectivity between populations corresponding to different potential crossing points across the Mwanza Gulf, based on water depth, crossing distance, and shoreline connectivity (Fig. 3). For each hypothesis, we computed shoreline-constrained distances between sites using the *geosphere* package (Hijmans *et al*., 2022). Morphological distances were obtained from PCA scores of dataset (2), to specifically test for isolation by distance along the parapatric gradient, and compared against geographical distances under each hypothesis using Maximum-Likelihood Population Effects (MLPE) models (Clarke *et al*., 2002), which account for non-independence due to shared sites and individual identity. Because the terminal Juma population may be ecologically displaced, we fitted each MLPE model on two datasets: one including all sites and one excluding Juma, to evaluate the influence of this population on spatial patterns.

**Figure 3.**
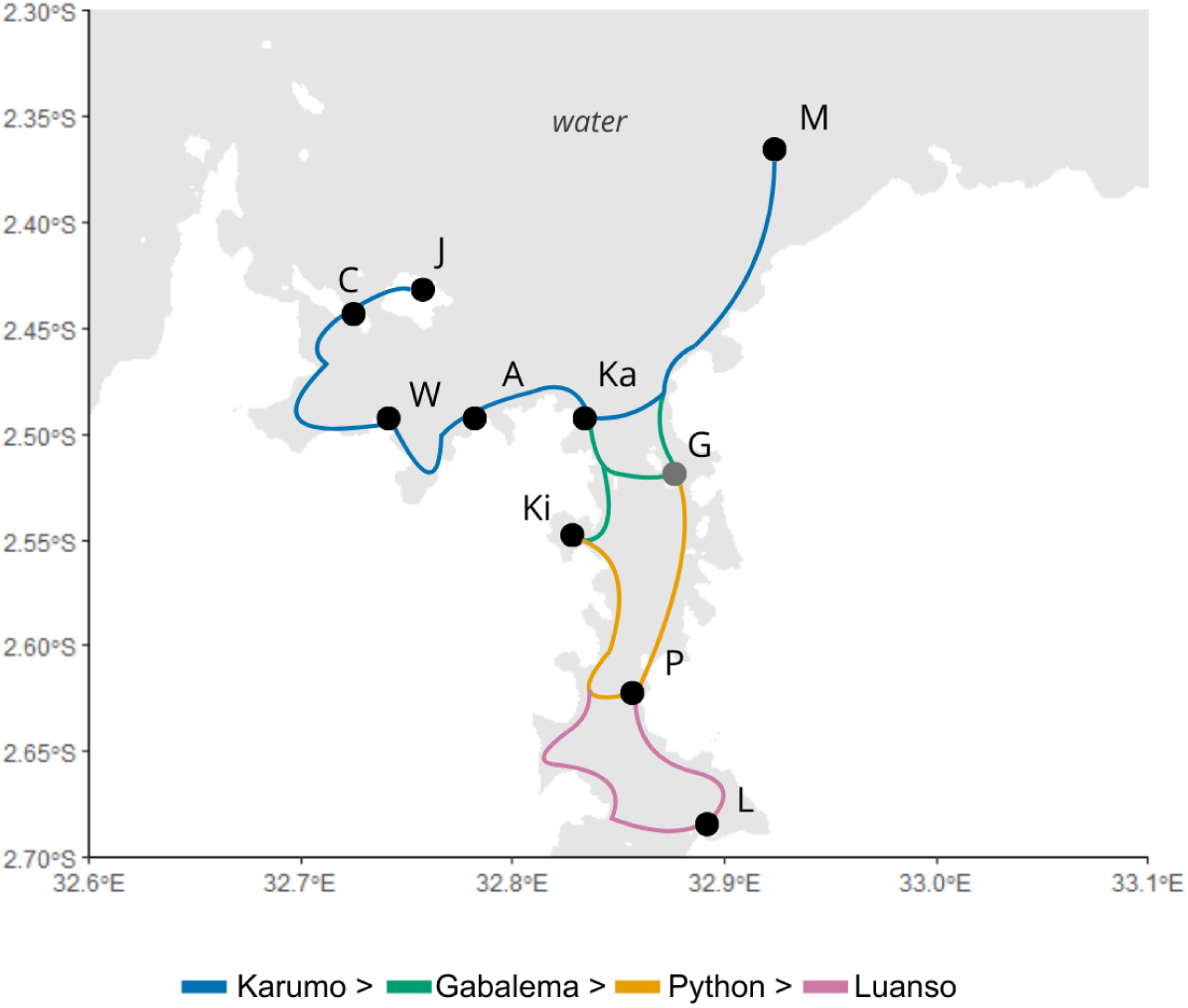
Spatial hypotheses of population connectivity and dispersal across the Mwanza Gulf for *Neochromis* and *Pundamilia* complexes. Four alternative shoreline-constrained dispersal pathways linking all sampled locations are shown in distinct colors, corresponding to different candidate crossing points across the gulf. In the Karumo and Gabalema alternatives, the southern locations (Python and Luanso) are connected following the east coast southwards Gabalema. In the Karumo alternative, Kissenda is linked following the west coast southwards Karumo. M: Makobe; G: Gabalema; P: Python; L: Luanso; Ki: Kissenda; Ka: Karumo; A: Amranda; W: Wakahangara; C: Chamagati; J: Juma.

We used bootstrap resampling (*n* = 1000) to derive 95% confidence intervals for the fixed effect of distance and compared spatial hypotheses using mean Akaike information criterion (AICc; Akaike, 1973; Burnham and Anderson, 2002; Hurvich and Tsai, 1993) values across bootstraps. To assess model robustness to local effects, we jackknifed locations and applied bootstrap resampling within each jackknife iteration (*n* = 1000) to obtain confidence intervals. The stability of slope estimates across jackknife iterations was quantified using the coefficient of variation (Brown, 1998), used as an index of model robustness. Finally, we estimated the relative effect of distance on craniofacial shape variation by multiplying the mean morphological dissimilarity between individuals by the bootstrapped fixed-effect coefficient from the best-supported model.

### Comparison of terminal species and ecological character displacement (ECD)

#### Multivariate approach

We used all PCA scores from dataset (3) to test for two key predictions of ECD: (i) greater interspecific divergence in sympatry vs. allopatry, and (ii) reduced intraspecific variance in sympatry (Anderson and Matute, 2025; Brown and Wilson, 1956). We performed unilateral permutation tests (*n* = 10,000) on multivariate pairwise distances for aforementioned predictions, with separate analyses for each species in prediction (ii). Bootstrap resampling (*n* = 10,000) was used to quantify uncertainty in our estimates and obtain confidence intervals for dissimilarity and variance under both conditions (sympatry/allopatry).

#### Trait-based approach

Trait convergence or directional shifts due to shared local environments can mask patterns consistent with ECD (Goldberg and Lande, 2006). To address this, we performed ANOVAs on individual traits (meeting parametric assumptions), testing for main effects and interactions of species and condition (sympatry/parapatry). We used a decision framework (Tab. 1) to interpret patterns based on combinations of main and interaction effects, as well as the direction and magnitude of trait shifts between sympatric and allopatric populations. This approach allows us to disentangle the contribution of ecological character displacement from local adaptation, distinguishing directional adaptations (traits increase or decrease in both species) from convergent adaptations (traits converge to similar values due to shared local conditions).

**Table 1.**
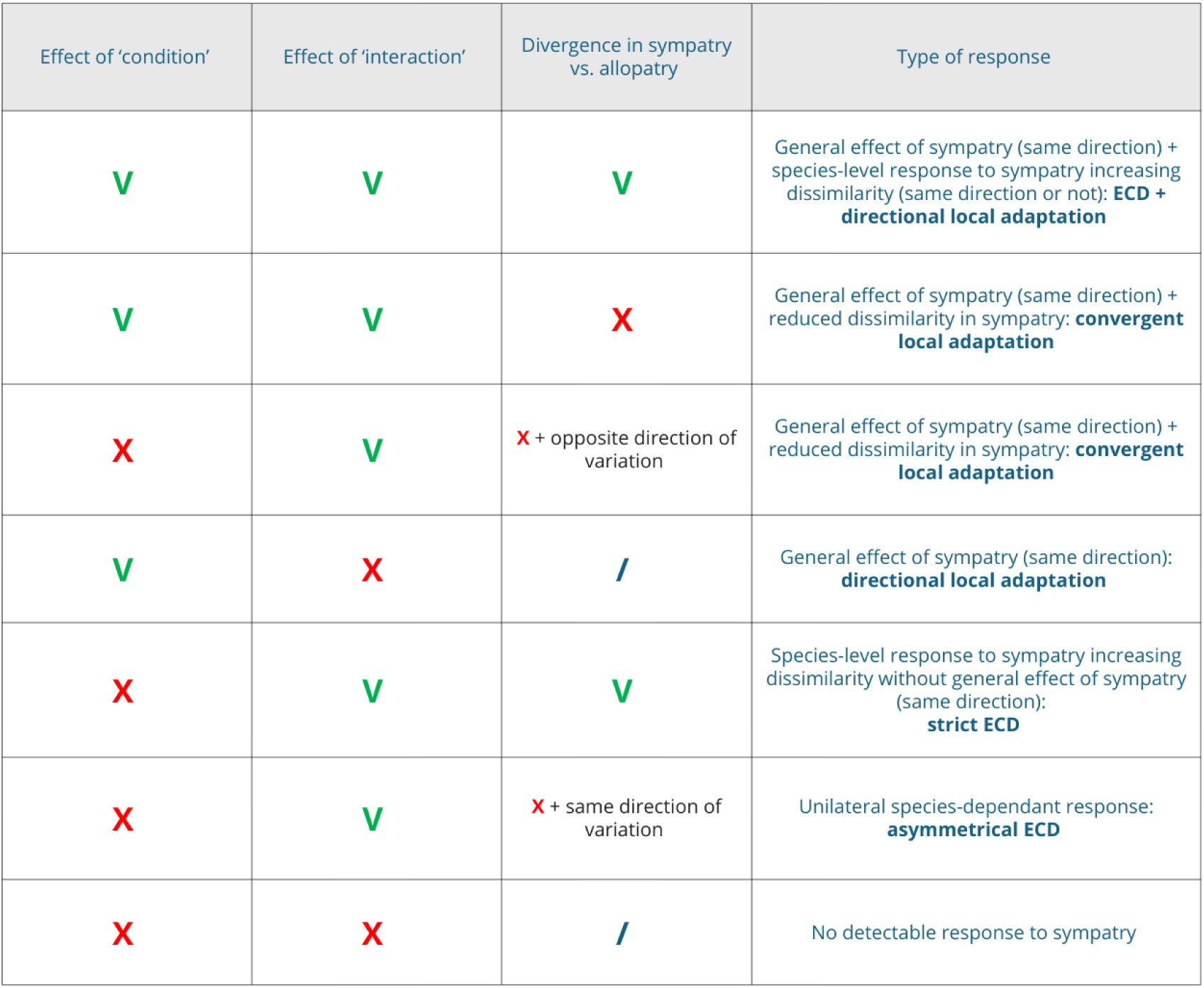
Decision framework for interpreting trait variation from ANOVAs. Significance of main effects of species and condition (sympatry/parapatry), as well as their interaction (species:condition), were combined with the directionality of mean trait shifts to distinguish alternative evolutionary scenarios (V: significant effect; X: non significant effect; /: not attributable).

| Effect of 'condition' | Effect of 'interaction' | Divergence in sympatry vs. allopatry | Type of response |
| --- | --- | --- | --- |
| V | V | V | General effect of sympatry (same direction) + species-level response to sympatry increasing dissimilarity (same direction or not): **ECD + directional local adaptation** |
| V | V | X | General effect of sympatry (same direction) + reduced dissimilarity in sympatry: **convergent local adaptation** |
| X | V | X + opposite direction of variation | General effect of sympatry (same direction) + reduced dissimilarity in sympatry: **convergent local adaptation** |
| V | X | / | General effect of sympatry (same direction): **directional local adaptation** |
| X | V | V | Species-level response to sympatry increasing dissimilarity without general effect of sympatry (same direction): **strict ECD** |
| X | V | X + same direction of variation | Unilateral species-dependant response: **asymmetrical ECD** |
| X | X | / | No detectable response to sympatry |

## RESULTS

### Morphological variation across distribution

#### Multivariate structure of craniofacial morphology

We examined overall patterns of craniofacial morphological variation in *Pundamilia* species complex (*n* = 111) and *Neochromis* species complex (*n* = 92) using Principal Component Analysis (PCA) on all individuals for each genus (dataset (1)), based on twelve measured traits (as defined in Materials and Methods).

In *Pundamilia*, the first four PC dimensions were associated with clear morphological domains. The first dimension (28.5%) includes traits associated with head and snout elongation as well as eye size. The second axis of variation (16.5%) relates to post-orbital robustness and head width. The third axis (12.3%) captures variation in eye size, and the fourth axis (10.7%) represents snout and lower jaw width. Trait clusters obtained through hierarchical clustering strictly match the morphological domains obtained with PCA loadings. These domains correspond to distinct and/or complementary functional modules—including head elongation, bite force, eye size, and mouth opening—known to predict feeding ecology in fish (Cooper and Westneat, 2009; Wainwright and Richard, 1995). In cichlids, these modules, and especially the preorbital components, have been regarded as major axes of anatomical variation across Great East-African lakes, with subsequent changes in trophic ecology across the clade’s evolution (Cooper *et al*., 2010).

In *Neochromis*, the first dimension accounts for a large part of the total variation (39.1%) with a strong positive contribution of all traits, indicating that the overall head size explains most of the craniofacial variation observed across the distribution. The second dimension (13.9%) contrasts head elongation to interorbital width, while the third (10.7%) contrasts cheek depth and body depth to eye size. These patterns may indicate allometric constraints or trade-offs, such as between bite force and vision for dimension 3. The fourth dimension (8.6%) primarily involves preorbital depth. The hierarchical clustering of traits revealed integrated trait groups similar to those in *Pundamilia*, but these groups did not correspond to the main PCA axes identified in *Neochromis*.

#### Spatial patterns of morphological variation

To visualize the spatial structure of morphological variation, we mapped the average PC scores per locality. In *Pundamilia*, we focused on PC1–PC4, which capture the main functional modules described above (Fig. 4a). Spatial structuring appeared at different scales depending on the functional module, but two main clusters emerged overall: the northwestern locations on one side (Juma, Chamagati, Wakahangara, Amranda), and the southern part of the Mwanza gulf together with the distant Makobe Island on the other. Neighbouring northwestern sites showed relatively similar values across functional modules, characterised by shorter, broader heads and wider mouths compared to the other cluster. However, local deviations disrupted these regional trends, with high PC3 scores (eye size) at Wakahangara and elevated PC4 scores (mouth width) in Juma Island *P.* sp. ‘big blue red’. Sympatric species on Juma Island exhibited strong morphological divergence, with scores clearly distinct from the geographically closest conspecific populations (Chamagati and Makobe). This spatial structure of PC scores is consistent with species distribution and highlights a sometimes gradual but also locally accentuated divergence in craniofacial shape.

**Figure 4a.**
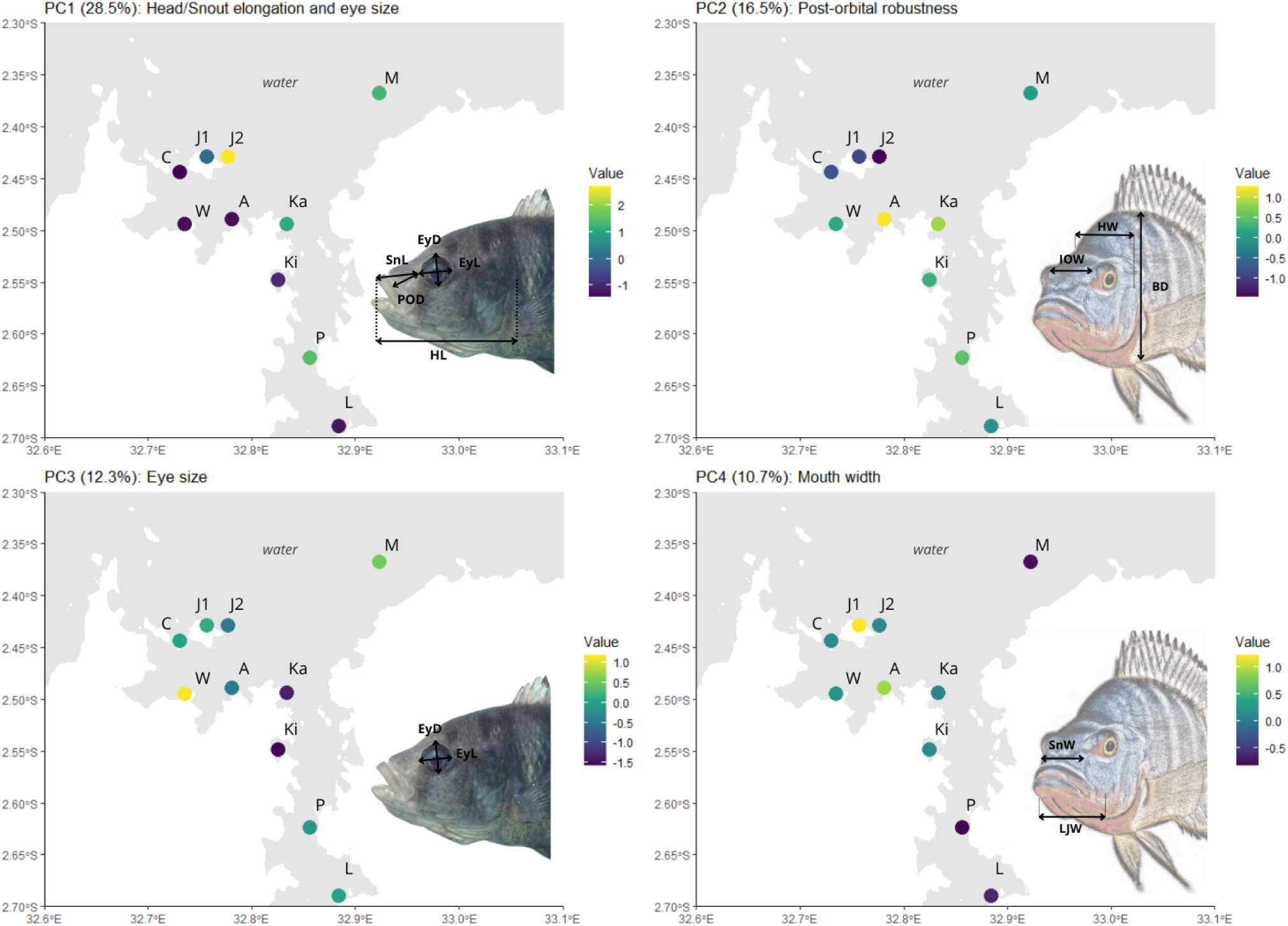
Heatmap of PC1-4 scores from the morphometric analysis of craniofacial shape across populations of *Pundamilia* (total *n* = 111). Traits with highest contribution are schematically illustrated for each PC, abbreviations are the same as in figure 2. On Juma island where species are sympatric, data is plotted separately for each species (J1, J2). M: Makobe; P: Python; L: Luanso; Ki: Kissenda; Ka: Karumo; A: Amranda; W: Wakahangara; C: Chamagati; J1: Juma *P*. sp. ‘big blue red’; J2: Juma *P*. *pundamilia*.

In *Neochromis*, we only mapped PC1 scores as a proxy of overall head size (Fig. 4b), as subsequent axes explained markedly less variation and did not correspond to clear functional modules, instead reflecting weaker allometric trade-offs with limited ecological interpretation (see Fig. S1 for PC1-PC4 representation). Extremes of head size were observed in Python (large) and Juma (small), while other locations showed relatively homogeneous values. This reduced variation limited the inference of spatial structuring. However, there is a clear divergence between sympatric species on Juma Island, revealing a pronounced differentiation of *N. omnicaeruleus* at this location compared to the conspecific population of Makobe.

**Figure 4b.**
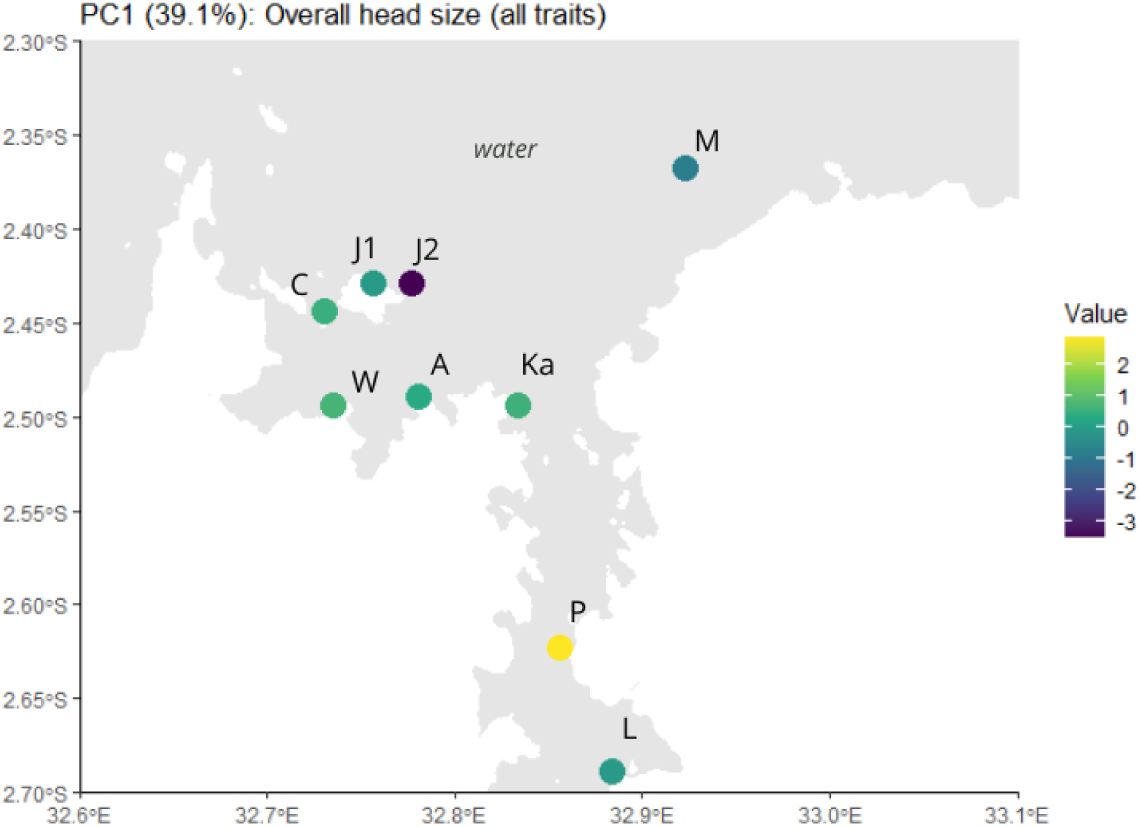
Heatmap of PC1 scores from the morphometric analysis of craniofacial shape across populations of *Neochromis* (total *n* = 92). On Juma island where species are sympatric, data is plotted separately for each species (J1, J2). M: Makobe; P: Python; L: Luanso; Ka: Karumo; A: Amranda; W: Wakahangara; C: Chamagati; J1: Juma *N. black nigricans*; J2: Juma *N. omnicaeruleus*.

Overall, craniofacial morphology in *Pundamilia* showed evidence of moderate spatial structuring, while both genera displayed clear morphological differentiation between sympatric species on Juma Island. These patterns motivated the subsequent analyses of isolation by distance (IBD) and ecological character displacement (ECD).

#### Isolation by distance

We applied a Maximum Likelihood Population Effect (MLPE) model on the multivariate scores obtained with PCA on the hypothesised parapatric gradient (dataset (2)) for each genus to test for the effect of IBD. In *Pundamilia*, all connectivity hypotheses produced positive distance–morphology relationships, both with and without the terminal population from Juma Island, and models with more northern crossing points across the Mwanza Gulf consistently yielded steeper IBD slopes (Fig. 5a). Among them, the Gabalema-crossing hypothesis was clearly the best-supported (ΔAICc = 26-41 with Juma;, 8–90 without), showing the lowest AICc value, including or excluding Juma’s population.

**Figure 5a.**
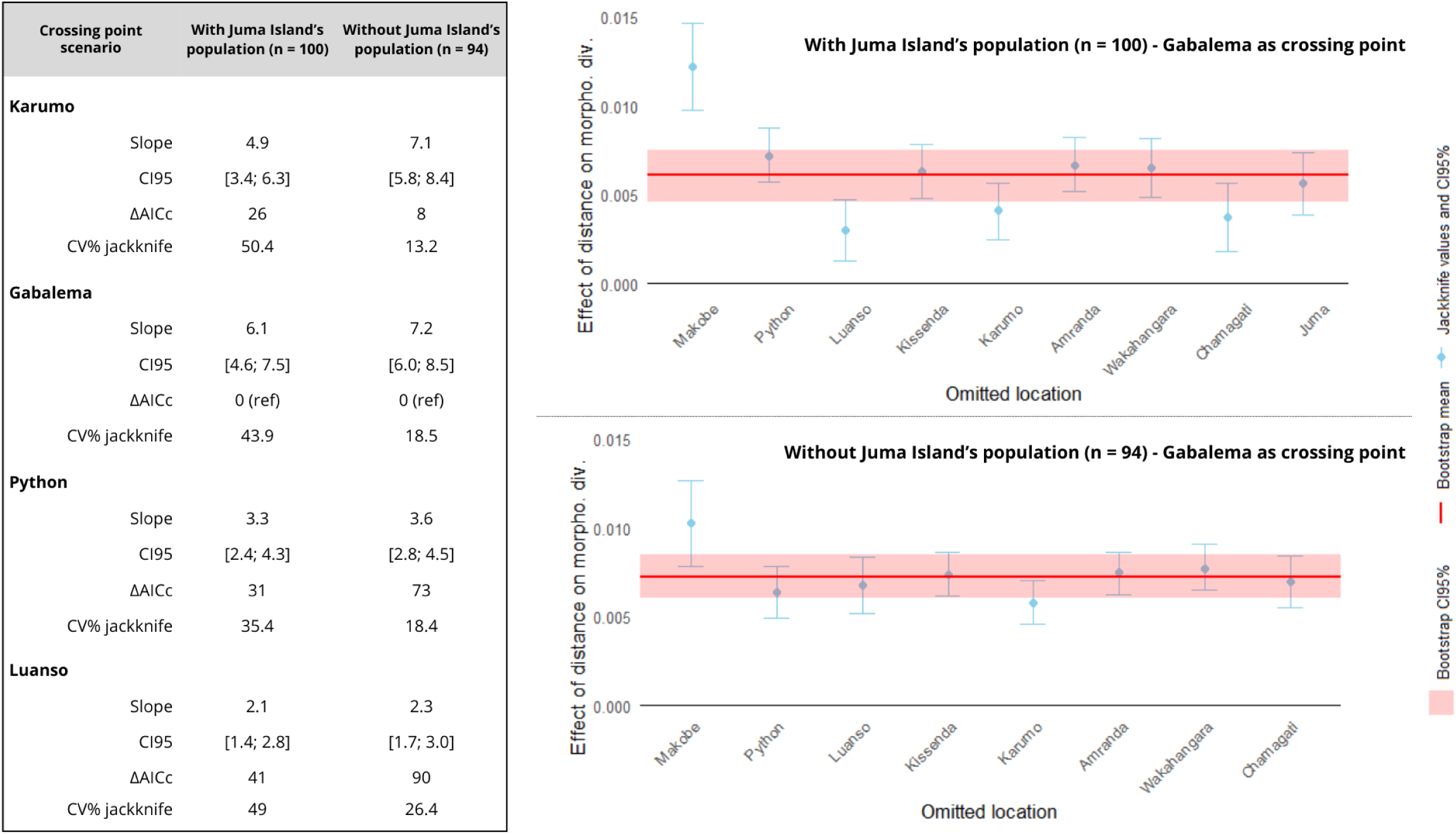
Isolation by distance analyses in *Pundamilia* under four alternative hypotheses of geographic connectivity, fitted using Maximum-Likelihood Population Effects (MLPE) models. The table summarises, for each connectivity scenario and for analyses performed with or without the terminal population at Juma Island, the MLPE slope (×10³), its 95% bootstrap confidence interval (×10³), the ΔAICc values, and the coefficient of variation of jackknife estimates (CV% jackknife). Across scenarios, removing the Juma population consistently increased model stability, reflected in lower CV% jackknife values. The Gabalema-crossing scenario, which had the strongest support (lowest AICc), is illustrated on the right panel, showing mean slopes, their 95% bootstrap confidence intervals, and jackknife diagnostics for analyses with and without Juma. See Fig. S2 for the representation of all hypotheses, with and without Juma Island’s population.

Removing the Juma population of *P.* sp. “big blue red” substantially improved model stability: jackknife variability, which could exceed 50% in the full dataset, declined across all hypotheses in the restricted dataset and dropped below 20% in three of four scenarios. The models’ robustness to single-location removal supports the absence of a dominant local influence on the global estimate of IBD, although individuals from Makobe Island tended to attenuate this estimate. The exclusion of Juma also increased the estimated strength of the IBD signal. Under the best-supported model, the predicted increase in craniofacial dissimilarity rose from 0.127% per km (≈12.5% between the most distant individuals) to 0.151% per km (≈14% maximum). Overall, the IBD pattern in *Pundamilia* is positive and the exclusion of the terminal Juma population reveals a consistent and robust clinal pattern, while the inclusion of the terminal population obscures and destabilises the underlying spatial signal.

In *Neochromis*, fitting the MLPE on the complete range resulted in weak or undetectable IBD (Fig. 5b). Across connectivity hypotheses, most 95% bootstrap intervals encompassed zero, and the best-supported model (a crossing point at Python Islands’ level) yielded a weak estimated effect, with a predicted increase in morphological dissimilarity of 0.029% per km (≈4.4% between the most distant individuals). The near-zero mean slope strongly inflated the CV of jackknife estimates, yet jackknife intervals remained highly regular across sites and connectivity hypotheses, except for the contribution of Juma, whose removal consistently shifted model behaviour towards higher estimates (Fig. S2).

**Figure 5b.**
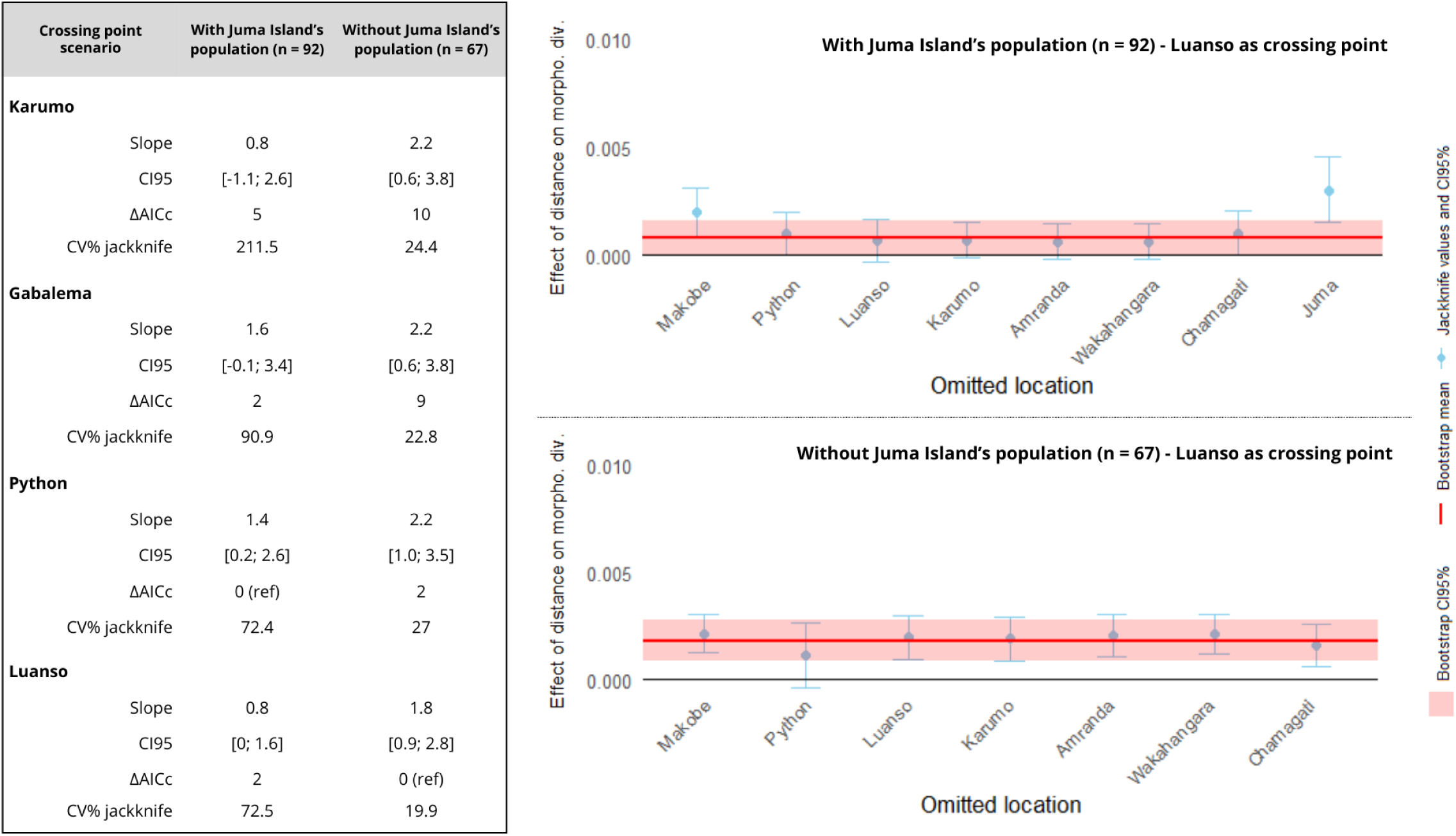
Isolation by distance analyses in *Neochromis* under four alternative hypotheses of geographic connectivity, fitted using Maximum-Likelihood Population Effects (MLPE) models. The table summarises, for each connectivity scenario and for analyses performed with or without the terminal population at Juma Island, the MLPE slope (×10³), its 95% bootstrap confidence interval (×10³), the ΔAICc values, and the coefficient of variation of jackknife estimates (CV% jackknife). Across scenarios, removing the Juma population consistently increased model stability, reflected in lower CV% jackknife values. The Luanso-crossing scenario, which had the strongest support after removing Juma (lowest AICc), is illustrated on the right panel, showing mean slopes, their 95% bootstrap confidence intervals, and jackknife diagnostics for analyses with and without Juma.See Fig. S2 for the representation of all hypotheses, with and without Juma Island’s population.

Excluding the population of *N. black nigricans* of Juma Island revealed a weak but robust signal of IBD. Under these settings, all spatial hypotheses produced positive and more stable IBD slopes, with jackknife CV values indicating robust estimates and little evidence for localised influence on the observed effect, except for Python Islands. The Luanso-crossing scenario received the strongest support (lowest AICc), with a slope corresponding to a predicted 0.038% increase in dissimilarity per km, corresponding to approximately 7% of craniofacial dissimilarity between both ends of the restricted range.

Across genera, the exclusion of the terminal Juma population strengthened and stabilised the IBD signal, marking its disproportionate local influence and highlighting the value of investigating ECD-related effects. Interestingly, *Pundamilia* showed a steeper IBD signal, with a spatial layout of differentiation best explained by connectivity across a northern crossing point of the Mwanza Gulf, whereas *Neochromis* expressed a weaker gradient with support for a more southern crossing. Together, these results reveal consistent but genus-specific patterns of clinal morphological differentiation along the parapatric gradient.

### Ecological character displacement

#### Multivariate approach

We assessed the predictions of ecological character displacement (ECD) using multivariate scores (dimensions 1 to 4) from PCA on dataset (3) for each genus. Specifically, we tested whether sympatric populations exhibit (i) greater interspecific divergence and (ii) reduced intraspecific variance, relative to allopatric populations.

For both genera, we found significantly greater interspecific morphological dissimilarity in sympatry, in line with ECD predictions (permutation test; *p* < 0.001; Fig. 6). In contrast, directed tests for intraspecific variance did not decrease in sympatry (*p* > 0.9), providing no support for this ECD prediction. Visual inspection of bootstrap distributions suggested a potentially opposite trend, with higher variance in sympatric populations. Exploratory post-hoc tests in the opposite direction supported this trend of a higher intraspecific variance in sympatry for *Pundamilia* (*P.* sp. big blue: *p* = 0.071; *P. pundamilia*: *p* = 0.022) and *Neochromis* (*N. black nigricans: p* = 0.068; *N. omnicaeruleus*: *p* = 0.042). The latter results should be interpreted cautiously, as they are not part of the a priori ECD predictions.

**Figure 6.**
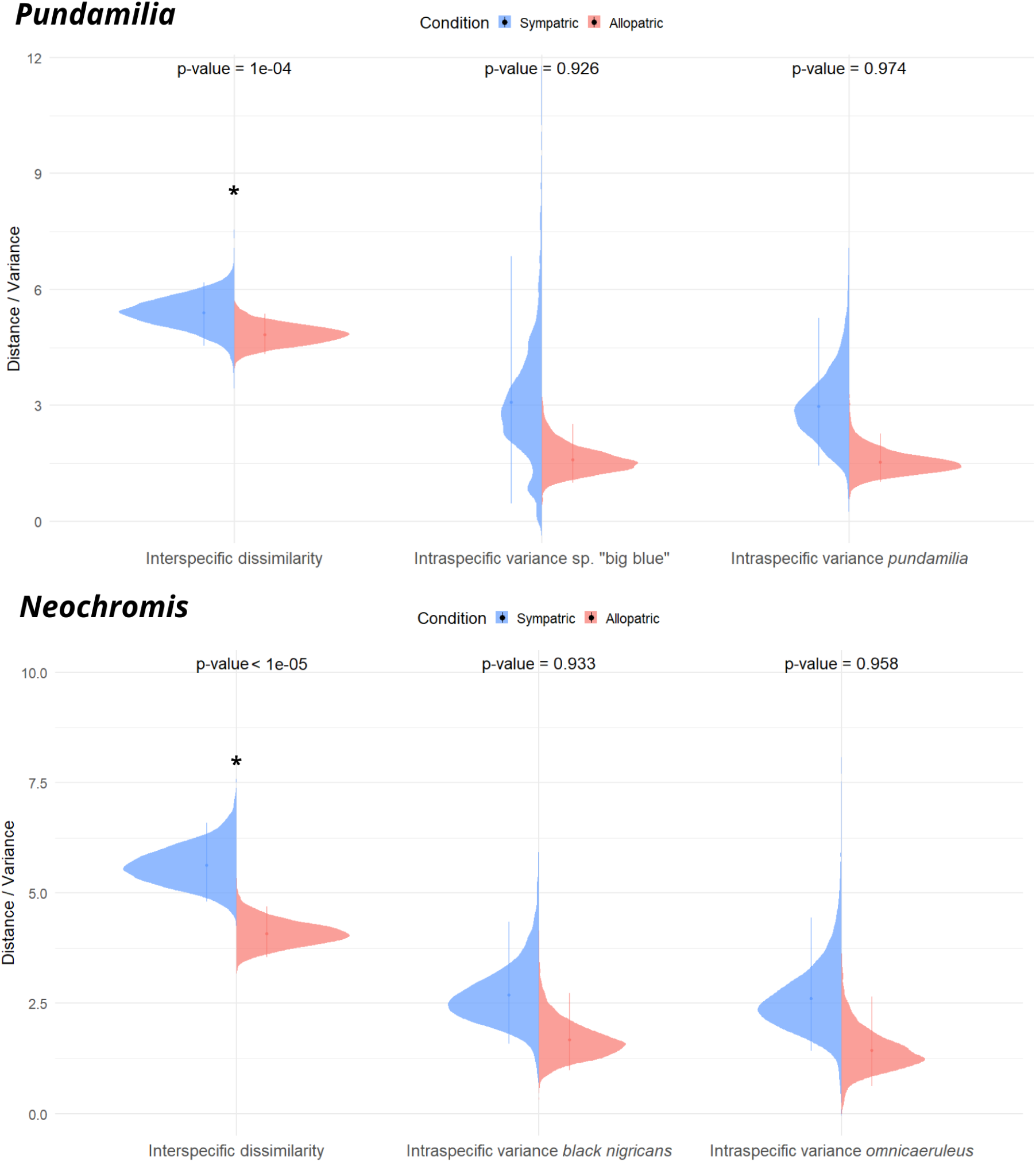
Comparison of interspecific dissimilarity and intraspecific variance in craniofacial morphology *Pundamilia* (top; *n* = 47) and *Neochromis* (bottom; *n* = 50) populations under sympatric versus allopatric conditions. Bootstrap distributions are shown for each comparison, with the same ordinate scale for dissimilarity and variance; vertical lines indicate the 95% CI. *P*-values displayed above each comparison correspond to permutation tests predicting greater interspecific dissimilarity and lower intraspecific variance in sympatry, according to ECD expectations.

#### Trait-based approach

The trait-based analysis of ECD using ANOVAs indicated that variation in craniofacial morphology reflects both ecological character displacement and local adaptation (Tab. 2; see Fig. S3 for boxplots of trait values). Effects were supported by statistically significant (p < 0.005) or near-significant differences (p < 0.1) in either the main effect of Condition (sympatric vs. allopatric) or the Condition x Species interaction.

**Table 2.** Classification of craniofacial traits according to evolutionary scenarios inferred from ANOVA in *Pundamilia* (*n* = 47) and *Neochromis* (*n* = 50). Categories include strict ecological character displacement (ECD), ECD combined with local adaptation (LA), convergent local adaptation, and directional local adaptation as described in Table 1. Reported p-values correspond to the effects of **Condition** (sympatric vs. allopatric) and the **Condition x Species** interaction. Traits shown in **bold** indicate statistically significant results supporting the inference (**p < 0.05**), while traits marked with an asterisk (*) represent near-significant values suggesting a potential effect (*0.05 ≤ p < 0.1*).

| Category | Genus | Trait | p (Condition) | p (Cond:Sp.) |
| --- | --- | --- | --- | --- |
| Strict ECD | *Pundamilia* | — | — | — |
|  | *Neochromis* | Lower jaw length | 0.497 | **0.036** |
| ECD + LA | *Pundamilia* | Body depth* | 0.056 | **0.037** |
|  |  | Cheek depth* | 0.055 | 0.061 |
|  | *Neochromis* | Eye length | **<0.001** | **<0.001** |
|  |  | Eye depth* | **0.002** | 0.058 |
| Convergent LA | *Pundamilia* | Interorbital width* | 0.644 | 0.052 |
|  | *Neochromis* | Cheek depth | **0.002** | **0.026** |
| Directional LA | *Pundamilia* | Head length | **0.002** | 0.950 |
|  |  | Snout width* | 0.078 | 0.318 |
|  |  | Eye length* | 0.068 | 0.241 |
|  |  | Preorbital depth | **<0.001** | 0.428 |
|  | *Neochromis* | Head length | **0.031** | 0.137 |
|  |  | Snout length | **0.039** | 0.355 |
|  |  | Snout width | **0.044** | 0.752 |

In *Pundamilia* (*n* = 47), sympatry was associated with enhanced divergence in body depth and cheek size, with *P. pundamilia* having shallower bodies and larger cheeks, and *P.* sp. ‘big blue red’ showing deeper bodies and smaller cheeks. Larger cheeks are consistent with greater adductor musculature and increased bite force (Cooper *et al*., 2011; DeLorenzo *et al*., 2022; Konings, 2016; Liem, 1973), whereas deeper bodies may improve maneuverability and stabilisation in structurally complex rocky habitats (Feilich, 2016). We also detected convergence in interorbital width, alongside a general elongation of the head. This elongation was characterised by longer heads, longer and broader snouts (snout length increased in both species, although not significantly), and longer eyes; traits previously linked to craniofacial functional performance (DeLorenzo *et al*., 2022; Konings, 2016).

In *Neochromis (n* = 50), sympatric populations showed greater divergence between species, with *N. black nigricans* exhibiting longer jaws and larger eyes, and *N. omnicaeruleus* showing shorter jaws and smaller eyes. Both species also displayed a general reduction in eye size consistent with local adaptation. Local environmental conditions on Juma Island appear to drive convergence in cheek depth, as both species exhibited reduced variance and similar mean values despite differing magnitudes of change. Additionally, individuals from Juma Island had shorter heads and smaller snouts than those from allopatric sites, independent of species identity.

## DISCUSSION

How variation in eco-morphological and sexual traits evolves across within a lineage to give rise to new and ecologically different species is not well understood. Our investigation into rocky shore cichlid fishes from Lake Victoria reveals complex lineage-specific patterns of IBD, local adaptation, and ECD shaping the evolution of species along a parapatric continuum. We propose that the observed patterns may potentially illustrate speciation in a geographic ring, but this hypothesis requires further study.

### Ecological and morphological divergence across geography

Our analysis of craniofacial variation across continuous geographic space found different patterns of craniofacial variation between the two complexes. In *Pundamilia*, the variation was structured around distinct functional modules, namely head and snout elongation, post-orbital robustness and width, eye size, and mouth width. These modules relate to diet-specific constraints. In particular, individuals with elongated head and snout and larger eyes are commonly associated with insectivory in cichlids, as better vision and quicker jaw movements enhance the capturing of mobile prey (DeLorenzo *et al*., 2022; Rüber and Adams, 2001). Herbivores, and especially algae grazers, display a wider mouth opening with strong bite force, permitted by shorter jaws and large muscles inserted in the post-orbital area (DeLorenzo *et al*., 2022; Westneat, 2003). The spatial structuring of module values highlights a progressive differentiation of craniofacial morphology in *Pundamilia* around the gulf, mirroring the gradual shift in trophic ecology (Seehausen, 1996) changing from insectivory in *P. pundamilia* to algae grazing in *P*. ‘big blue red’ (Fig. 1a).

Analyses of isolation by distance (IBD) also revealed a robust and moderate positive effect of geographic distance on morphological dissimilarity across the parapatric range (estimated at ∼1.5% every 10km). Notably, the population of *P.* sp. “big blue red” from Juma Island produced a destabilising effect on the analysis, most likely attributable to local effects, such as ECD, which are discussed below. Its exclusion greatly improved the quality of the signal. Individuals from Makobe also tended to attenuate the IBD slope, suggesting a disproportionate local influence on the global pattern. This effect may reflect the geographic isolation of Makobe Island, located 5km offshore and surrounded by deep waters, potentially imposing stronger resistance to dispersal than expected from linear shoreline distance alone. Model comparison unequivocally favoured a crossing point at Gabalema, in the northern part of the Mwanza Gulf, yet all connectivity hypotheses yielded positive and stable IBD slopes. This pattern may reflect the presence of multiple, partially overlapping spatial patterns of evolution. In particular, the northern part of the gulf may have provided older and more persistent connections during the lake’s refilling (14-10kya; Wienhues *et al*., 2024), whereas southern crossing events may have been more recent or less frequent. This suggests that population dispersal along rocky shorelines in *Pundamilia* may have taken multiple, variable routes over time across the gulf.

Environmental gradients such as water depth, turbidity, or rock size and shape, as well as interspecific interactions, directly shape trophic niches and prey availability in Lake Victoria’s species-rich waters (Conith *et al*., 2020; Ford *et al*., 2016; Seehausen and Bouton, 1997). Consequently, the signal of IBD captures both the cumulative effects of gradual divergence along the continuous geographic gradient and the influence of local effects, providing insight into the interplay between historical processes and more recent local adaptation.

The gradual eco-morphological divergence in *Pundamilia* is paralleled by shifts in male nuptial coloration, ranging from blue-grey to bright sky-blue (Fig. 1a). Turbidity is unlikely to explain these changes, as assortative mating in turbid waters primarily relies on red tones (Carleton *et al*., 2005), and water transparency is similar across locations with different colour morphs (Seehausen, 2006). However, many sites host multiple cichlid species, and shifts in male coloration may partly reflect selection to enhance species discrimination in these multispecies communities. Such interspecific effects can interact with assortative mating within populations, reinforcing both coloration divergence and ecological differentiation (Couldridge and Alexander, 2002; Seehausen and Schluter, 2004). Additionally, accumulated differences via assortative mating likely strengthen reproductive isolation, potentially acting in concert with IBD to maintain divergence despite ongoing gene flow (Ding *et al*., 2014; Salzburger, 2009; Seehausen *et al*., 1998). Together, these patterns suggest that ecological (craniofacial morphology) and sexual traits in *Pundamilia* reflect the combined influence of long-term evolutionary history, local adaptive events, and ecological interactions, consistent with parapatric speciation along a spatially structured gradient.

In *Neochromis*, craniofacial variation was largely explained by overall head size rather than discrete functional modules, with PC1 alone accounting for nearly 40% of variance. This pattern suggests a lower degree of morphological modularity compared to *Pundamilia*, and may reflect stronger allometric constraints or a reduced range of trophic specialisation in these predominantly generalist–herbivorous species (Hulsey and León, 2005; Martinez *et al*., 2018). Accordingly, the IBD pattern was weaker, consistent with the idea that strong allometric integration and limited trophic specialisation constrain the extent to which ecological gradients and distance can generate divergence (Rüber *et al*., 1999; Voje *et al*., 2014). Nevertheless, MLPE analyses revealed a weak but robust effect of distance on morphological divergence. As in *Pundamilia*, the population of *N. black nigricans* from Juma Island produced a destabilising influence on the IBD signal, despite not exhibiting extreme PC1 scores. This suggests that interactions with the sympatric *N. omnicaeruleus*—characterised by markedly smaller heads— could represent a case of ECD. The best-supported models favoured hypotheses where connectivity integrates crossing points in the southern part of the Mwanza Gulf. These southern-crossing scenarios paralleled the gradual changes in male nuptial coloration described by Seehausen (1996), suggesting that both craniofacial and sexual traits follow a shared geographic gradient of divergence. Given that the more Southern parts of the Mwanza gulf filled later in time (Wienhues *et al*., 2024), it is possible that the *Neochromis* populations dispersed along the shoreline after the *Pundamilia* populations. Superimposed on this background divergence, localised effects disrupted the IBD signal in *Neochromis*. At Python Islands, unusually large head sizes may reflect introgression with *Mbipia mbipi* (Seehausen, 1996), producing outlier values that locally inflate morphological divergence.

Overall, both complexes show gradual geographic shifts in craniofacial morphology accompanied by changes in male coloration, consistent with clinal parapatric divergence. Yet, while these patterns in *Pundamilia* are closely tied to functional trophic specialisation, craniofacial divergence in *Neochromis* appears weaker and more strongly modulated by localised ecological or potential hybridisation effects.

### Ecological and sexual character displacement

The multivariate analysis on focus species’ morphology revealed an increase in interspecific dissimilarity in sympatry, relative to allopatry, in both genera, as expected in ECD settings (Brown and Wilson, 1956). In *Pundamilia*, this divergence in sympatry was particularly marked for cheek and body depth, and echoed species’ trophic ecology: the insectivorous *P. pundamilia* displayed enlarged cheeks, associated with a stronger bite force (Cooper *et al*., 2011; DeLorenzo *et al*., 2022; Konings, 2016), and shallower bodies, conferring enhanced predation success on benthic and shelled invertebrates (Hulsey and León, 2005; Liem, 1991) in rock crevices at <0.5m water depth where it typically forages (Seehausen, 1996). By contrast, the algae-scraping *P.* sp. ‘big blue red’ that is only found at depths of 0.5 - 4.0m, displayed less developed mandibular features but deeper bodies, the latter being essential for stability and manoeuvrability during foraging in algivorous cichlids (Barel, 1982; Feilich, 2016). This increased divergence at the sympatric location likely reflects ecological specialisation and depth-associated niche partitioning, reducing interspecific competition (Bouton *et al*., 1997).

Sympatry in *Pundamilia* was also associated with convergent and directional changes, including a general elongation of the head and intermediate values of interorbital width, which may be explained by environmental constraints such as the size of rock boulders. Such a combination of local adaptation and ECD has been documented in other competing species (Vadas, 1990), highlighting the interplay of multiple evolutionary forces in maintaining species coexistence.

In *Neochromis*, sympatric species showed increased divergence in lower jaw length and eye size, despite having similar diets. This pattern could indicate interspecific territoriality, a form of spatial micro-distribution often observed in male cichlids where territoriality and aggression maintain ecological segregation between species with overlapping trophic niches (Bouton *et al*., 1997; Takamura, 1984; Vadas, 1990). In our case, although depth ranges overlap between the two species on Juma (0.5 - 4m in *N. black nigricans*, and down to 6m in *N. omnicaeruleus*; Seehausen, 1996) the longer jaws and eyes of *N. black nigricans* may provide greater acuity and agility in narrower patches and crevices, while *N. omnicaeruleus* may forage more on gentle boulders and open micro-habitats.

Interestingly, morphological changes attributed to local adaptation in *Neochromis* were opposite to those observed in *Pundamilia*, with both *Neochromis* species showing shorter heads, smaller eyes and smaller snouts. It remains unclear whether these contrasting patterns reflect differences in evolutionary history, in niche partitioning, or in ecological interactions with the numerous species inhabiting Juma Island.

The multivariate analysis also revealed an increase in intraspecific variance in sympatry in both genera, contrary to ECD predictions. In *P.* sp. ‘big blue red’ (the sympatric morph of *P.* sp. ‘big blue’), the variance does not appear normally distributed (Fig. 5), and may reflect an intraspecific partitioning of individuals, as Seehausen (1996) indicates that “the very big fully adult males are restricted to the deeper parts of the species’ depth range and can easily be taken for a separate species”. This ecological stratification of large individuals undoubtedly influences morphology (Conith *et al*., 2020; Magalhaes *et al*., 2009), and could be the major source of intraspecific variance in *P.* sp. ‘big blue red’. More generally, three factors could contribute to elevated variance at Juma island. First, the species-rich community favours introgression and hybridisation, processes known to increase genetic and phenotypic variability (Lucek *et al*., 2017). Second, strong environmental heterogeneity, as observed on Juma Island, and habitat fluctuations can promote phenotypic plasticity and thereby morphological variance (Chevin and Chauhan, 2025; Magalhaes *et al*., 2009; Parsons *et al*., 2016). Third, strong sexual selection on coloration may relax ecological constraints on trophic morphology. While still insufficiently tested, this idea is consistent with evidence that divergent sexual selection on coloration promoted diversification after trophic adaptation in Lake Victoria, with assortative mating playing a central role in maintaining segregation among closely related species (Danley and Kocher, 2001; Doorn *et al*., 1998; Salzburger, 2009; Seehausen *et al*., 1998).

Finally, both genera also display important sexual character displacement on male nuptial coloration. The abrupt change from bright blue to reddish coloration between Chamagati and Juma likely increases assortative mating and isolation between sympatric *P.* sp. ‘big blue red’ and *P. pundamilia* (Maan *et al*., 2004; Seehausen, 1997; Svensson *et al*., 2024). Similarly, the shift to a completely black morph in *N. black nigricans* likely strengthens the divergent sexual selection that arises in allopatry between *N. omnicaeruleus and N. greenwoodi* (Magalhaes *et al*., 2010). These sexual character displacements act in concert with ECD to maintain species coexistence: the former maintaining reproductive isolation (Kopp *et al*., 2018; Seehausen *et al*., 1998), and the latter reducing competitive exclusion and preventing extinction (Doorn *et al*., 1998).

### Evolutionary origin of sympatric species: towards a ring species scenario?

Understanding the evolutionary origin of the sympatric species pairs in our system is essential for interpreting the evolutionary processes that now allow their coexistence. Our analysis of IBD suggests that Juma island’s populations of *P.* sp. ‘big blue red’ and *N. black nigricans* represent the terminal ends of two parapatric population chains of *Pundamilia* and *Neochromis*, respectively, which display gradual ecomorphological and male nuptial colour variation along rocky shores of Southern Lake Victoria. The combination of ecological and sexual selection likely maintains a parapatric speciation continuum between Makobe and Juma Island, producing distinct terminal species despite ongoing gene flow (Cooney *et al*., 2017; van Rijssel *et al*., 2018).

The other sympatric populations at the end of the cline at Juma island, *P. pundamilia* and *N. omnicaeruleus*, strongly resemble the conspecific populations found on Makobe Island, albeit with signs of local adaptation. Two evolutionary scenarios may account for their presence on Juma Island. Under the first scenario, these species originated through sympatric divergence on Juma. Ancestral populations could have experienced disruptive ecological selection acting on standing variation in morphology and genetically linked coloration, generating two distinct ecomorphs (Doorn *et al*., 1998; Kusche *et al*., 2015; Smith, 1966). In this case, the resemblance to Makobe taxa would reflect parallel or convergent evolution under similar ecological conditions (Muschick *et al*., 2012). Theory predicts that such divergence is facilitated on environmental gradients when spatially restricted dispersal and assortative mating generate locally disruptive, frequency-dependent selection (Doebeli and Dieckmann, 2000; 2003). These models show that evolutionary branching can occur within a continuous population distributed across a gradient, producing ecologically and phenotypically distinct lineages capable of coexisting in sympatry. Consistent with this interpretation, documented analyses of genetic differentiation at neutral loci in the sister species *P. pundamilia* (blue morph) and *P. nyererei* (red morph) provided strong support for repeated events of sympatric speciation, stemming from hybrid populations, within this lineage in the same geographical area (Meier *et al*., 2016; Seehausen *et al*., 2008).

The alternative scenario involves secondary contact following migration from Makobe or another nearby island. In this “secondary sympatry” model, coexistence is promoted by ecological and sexual divergence accumulated in allopatry (“species sorting”; Anderson and Matute, 2025), and later accentuated by character displacement (Anderson and Matute, 2025; Servedio, 2016). A particular variant of this scenario arises when secondary sympatry is maintained by recurrent migrations between Makobe and Juma, a process documented in Lake Victoria cichlids (Magalhaes *et al*., 2010; Terai *et al*., 2006). Under this assumption, the sympatric *P. pundamilia* and/or *N. omnicaeruleus* would be part of the chain of parapatric populations along the Mwanza gulf, thereby establishing a speciation continuum with gene flow between the apparently reproductively isolated sympatric species. This configuration is consistent with a ring-species process, in which a pair of sympatric species is linked by a chain of intergrading populations around a geographic barrier - deep water in this case (Irwin *et al*., 2005, 2001; Irwin and Wake, 2016; Mayr, 1963). Documented cases of ring speciation generally involve large spatial scales and long evolutionary times, and many fail to demonstrate continuous gene flow along the ring (Highton, 1998; Irwin and Wake, 2016; Liebers *et al*., 2004; Monjaraz-Ruedas *et al*., 2024). By contrast, in our evolutionarily young study system, geographical distances ( < 150 km) and divergence time ( < 16,000 years) are remarkably low, providing a promising framework to evaluate Mayr’s criteria (Martins *et al*., 2013; Mayr, 1963): (1) the presence of two terminal, sympatric populations that are reproductively isolated from one another; (2) each of the other populations that make up the ring must be currently, and have been historically, connected to their nearest neighbours through gene flow and this pattern must be the result of range expansion around a barrier; (3) all populations must be geographically connected forming a complete ring, and (4) the two terminal forms must be connected by gradual differences around the ring. The results obtained in this study only corroborate criterion (4) and to some extent criteria (1) -, but call for further genetic analyses and mate preference experiments to formally test for criteria (1)-(3) in both genera. If all criteria were to be met, this would be the first case of a ring species in freshwater fish, and provide valuable insights into the evolutionary forces that have driven the adaptive radiation of Lake Victoria cichlids.

## CONCLUSION

Our results reveal that isolation by distance (IBD) and local adaptation (LA) jointly shape ecomorphological divergence within parallel biogeographical clines of cichlid species. Coexistence of sympatric species at one end of the cline is promoted by phenotypic divergence of sympatric species, further reinforced by sexual and ecological character displacement (ECD). These findings highlight that speciation in young adaptive radiations is governed by the complex interplay of ecological divergence, sexual selection, and historical biogeography, with the relative contribution of each process varying across lineages. Future genomic analyses will be key to disentangle patterns of gene flow and to identify potential ring species in *Pundamilia* and *Neochromis*.

## ACKNOWLEDGEMENTS

We thank Salome Mwaiko (EAWAG) and Marcel Haesler (Uni. of Bern/EAWAG) for their help with the management of the fish collection. G. Gautier acknowledges the help of the Swiss-European Mobility Programme in providing financial support for his traineeship, the University of Bern and the EAWAG research centre for hosting the project, and the support of Prof. Jacintha Ellers and the Vrije Universiteit of Amsterdam for the traineeship supervision.

## AUTHOR CONTRIBUTIONS

Grégoire Gautier: Linear measurements, morphological and statistical analysis, and manuscript writing.

Anna Mahulu: Sampling, linear measurements methods, and contribution to writing. Ole Seehausen: Conception, sampling, and contribution to writing.

Pooja Singh: Conception, supervision, sampling, analysis, and contribution to writing.

## DATA AVAILABILITY STATEMENT

The data underlying this article are available in the article and in its online supplementary material.

## SUPPLEMENTARY FIGURES

**Figure S1.**
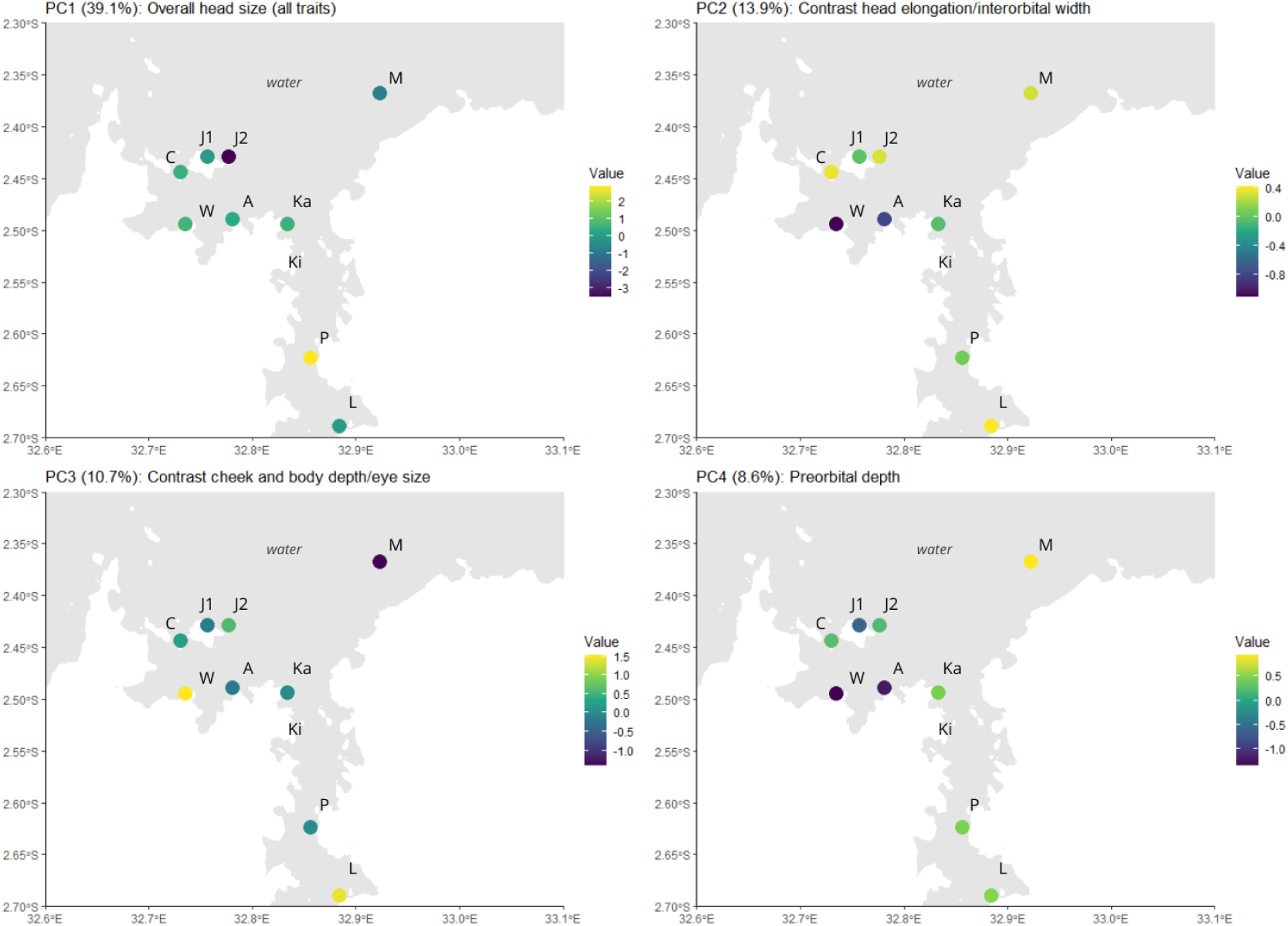
Heatmap of PC1-4 scores across locations in *Neochromis* (*n* = 92). Distinction between species was made on Juma Island. M: Makobe; P: Python; L: Luanso; Ka: Karumo; A: Amranda; W: Wakahangara; C: Chamagati; J1: Juma *N. black nigricans*; J2: Juma *N. omnicaeruleus*.

**Figure S2.**
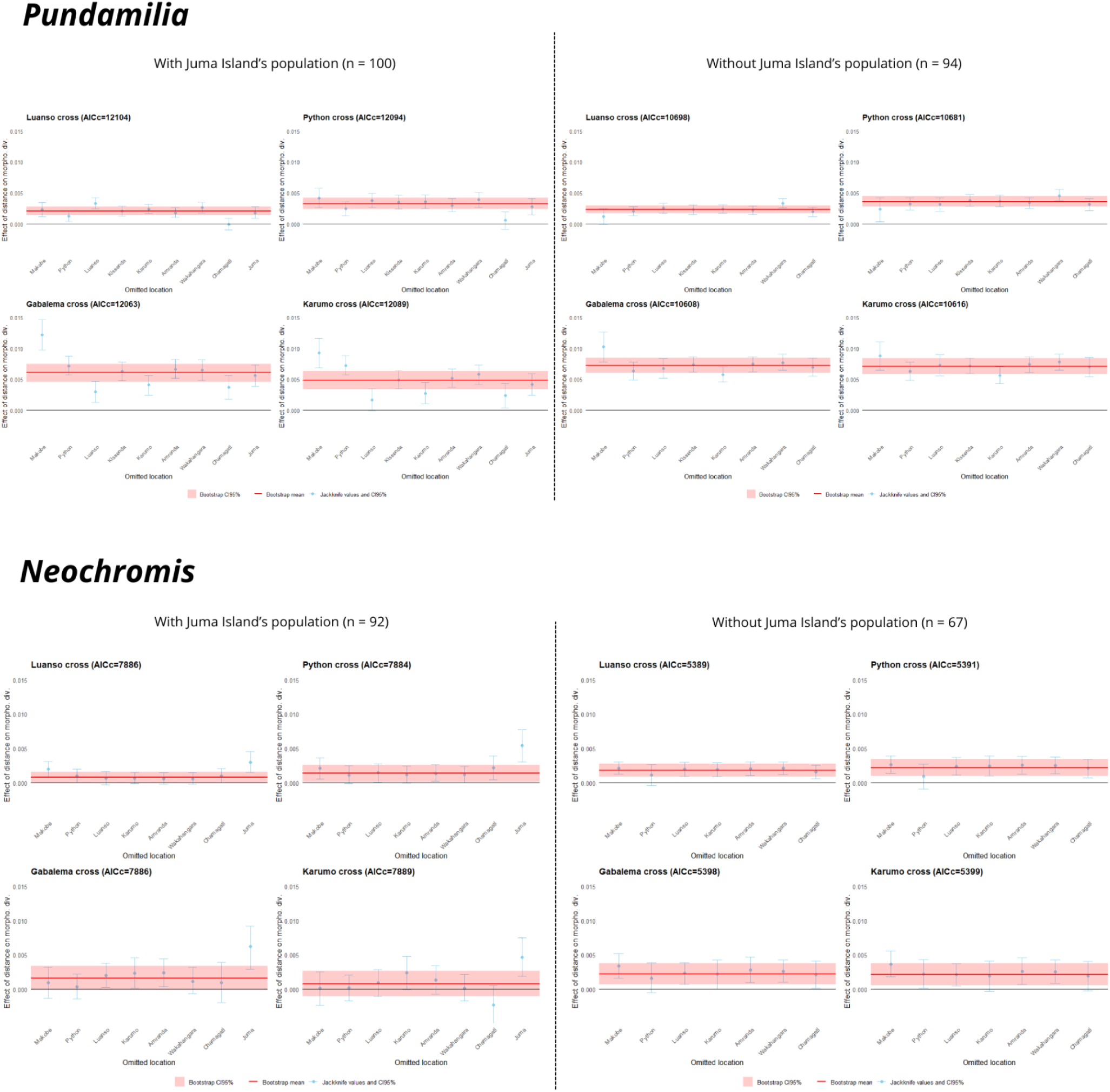
Results of the Maximum Likelihood Population Effect model (MLPE) analysis of isolation by distance in *Pundamilia* and *Neochromis*, across connectivity hypotheses, with and without Juma Island individuals. For each hypothesis, the mean slope estimate is shown with 95% bootstrap confidence intervals and jackknife diagnostics. Estimates and robustness indexes are inscribed in the panels of Fig. 4.

**Figure S3.**
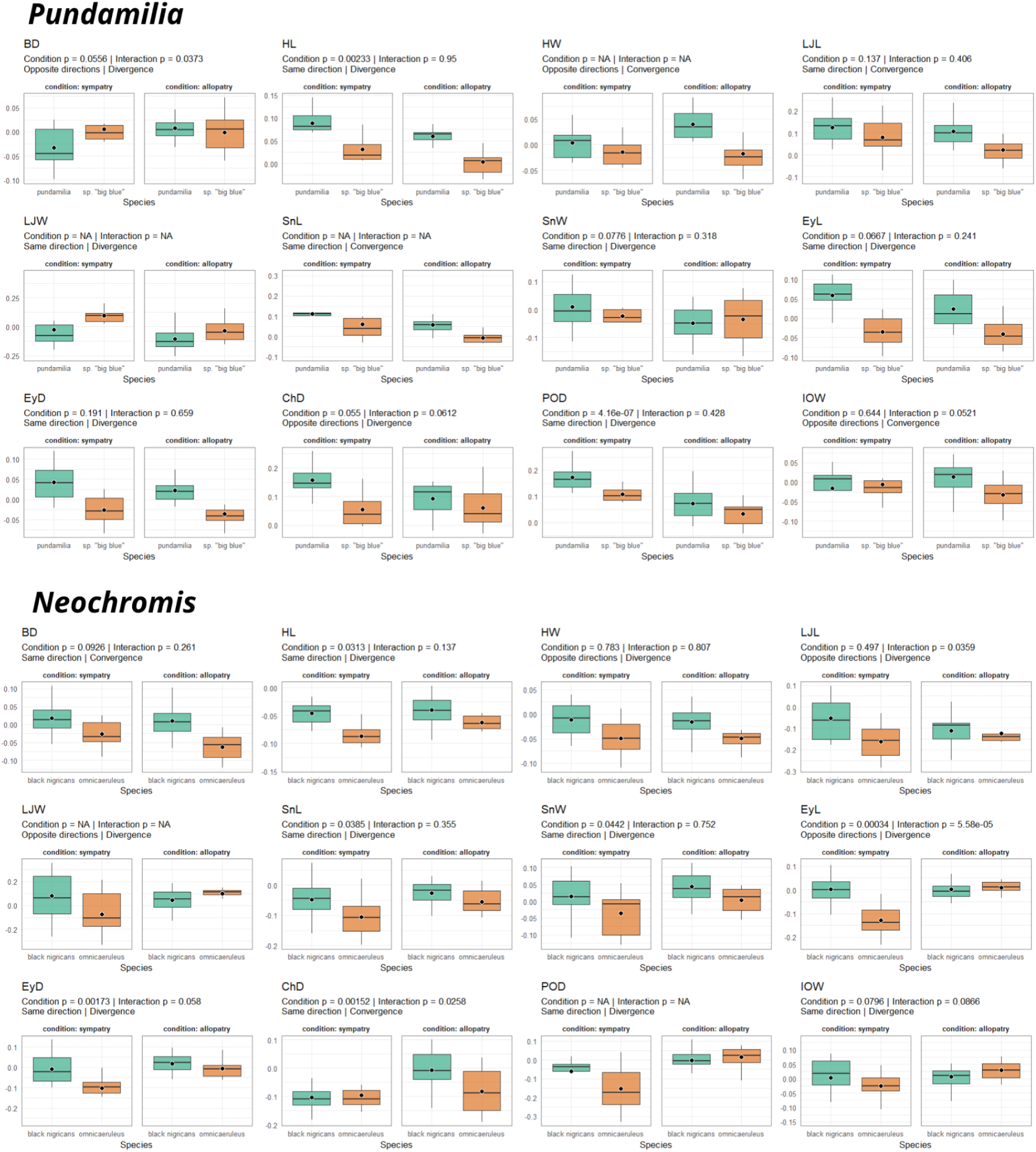
Boxplots of trait values across species and conditions, for ecological character displacement (ECD) analysis for *Pundamilia* and *Neochromis* lineages. We test ECD in allopatric versus sympatric populations for each morphological trait. *P*-values from the ANOVAs are indicated for the effect of condition (sympatry/allopatry) and the interaction species:condition. The direction and convergence/divergence of changes in trait values are indicated. BD: Body depth; HL: Head length; EyL: Eye length; EyD: Eye depth; ChD: Cheek depth; POD: Pre-orbital depth; SnL: Snout length; LJL: Lower jaw length; SnW: Snout width; LJW: Lower jaw width; IOW: Inter-orbital width; HW: Head width. X-axis lists only the species (not the genus).

